# HABIT – a webserver for interactive T cell neoepitope discovery

**DOI:** 10.1101/535716

**Authors:** Joana Martins, Carlos Magalhães, Vítor Vieira, Miguel Rocha, Nuno S. Osório

## Abstract

Neoepitopes generated by amino acid variants specifically found in pathogens or cancer cells are gaining momentum in immunotherapy development. HABIT (HLA Binding InTelligence) is a web platform designed to generate and analyse machine learning-based T cell epitope predictions for improved neoepitope discovery.

**Availability and Implementation:** HABIT is available at http://habit.evobiomed.com. Peptide-HLA binding prediction software were implemented in a web application for interactive data exploration using shiny package powered by RStudio.

**Supplementary information:** Supplementary data and video tutorials are available online.

## 1 Introduction

Immune responses mediated by T cells are triggered by specific peptides (T cell epitopes) bound to the highly diverse human leukocyte antigens (HLA) (Blum *et al.*, 2013). Variations in one single amino acid might impact peptide-HLA interactions and subsequent T cell responses (Unanue *et al.*, 2016; Hilf *et al.*, 2018). Many computational methods have been developed for predicting peptide-HLA binding improving the efficiency of T cell epitope identification (Zhang *et al.*, 2011). However, using this type of tools to identify neoepitopes, such as the ones generated by cancer or pathogen-specific amino acid mutations, is not trivial. It requires individual prediction of wild-type and mutant sequences in each tool, the selection of all overlapping peptides including the amino acid positions of interest and the comparison of the results generated separately. Thus, programming skills are needed to extract meaningful conclusions from the large outputs of HLA-peptide binding prediction data. Furthermore, data visualization and prioritization strategies, such as statistical tests or human population coverage calculation are inexistent in an integrated manner and would be of great value for immune-based therapy and diagnostic strategies (Oyarzun *et al.*, 2015). Taking this into account, we developed HABIT, which automates HLA binding prediction and provides advanced interpretation of the impact of amino acid variants in peptide-HLA binding (class I and II). It features an intuitive, user-friendly and interactive graphical interface that plots, summarizes and performs statistical comparisons on the antigenicity scores of wild-type and mutant epitope candidates.

## 2 Implementation

HABIT was developed in the RStudio v3.5.0, using the shiny v. 1.2.0 package (Chang *et al.*, 2018) to create the web-based application. Plotly v.4.8.0 (Sievert, 2018) and DataTables v.0.4 (Xiu *et al.*, 2018) packages were used for the generation of interactive plots and customizable tables. Peptide-HLA binding predictions tools (O’Donnell *et al.*, 2018; Zhang *et al.*, 2012) for HLA molecules were implemented in the back-end of HABIT to predict the peptide fragments with high binding affinity to HLA class I and class II molecules. All 8 to 14-mer peptides for wild-type and mutant sequences overlapping the position of the mutation of interest can be predicted to a set of class I alleles. Simultaneously, the binding affinity of 9-30 mer of peptides to HLA class II alleles can be predicted. Predicted antigenic scores are plotted and summarized enabling efficient selection of neoepitope candidates.

## 3 Results

To illustrate the relevance of the features implemented in HABIT, we analysed the effect of a somatic mutation (TP53 R248W) found in several types of human cancer (Tate *et al.*, 2018). Simultaneous HLA-binding prediction was launched for all wild-type and mutant peptides containing the amino acid residue of interest (Figure 1A and B). An interactive visual representation of the impact the R248W mutation had on HLA binding affinity was obtained. As shown in Figure 1C, the mutant amino acid generated several novel peptides predicted to bind strongly (in orange) to specific HLA class II proteins (Figure 1C). Interactive tables detailing the differences between wild-type and mutant peptides were also generated (Figure 1D), including percentage of strong binders, mean HLA affinity and mean percentile rank, along with statistical analysis. Sort and search functionality highlighted the HLA proteins with higher affinity for the majority of the neoepitopes (Figure 1D). Information about HLA distribution across the globe, as a way to evaluate the impact of neoepitopes in the immune responses across distinct human populations was also obtained (Figure 1E).

**Figure 1.**
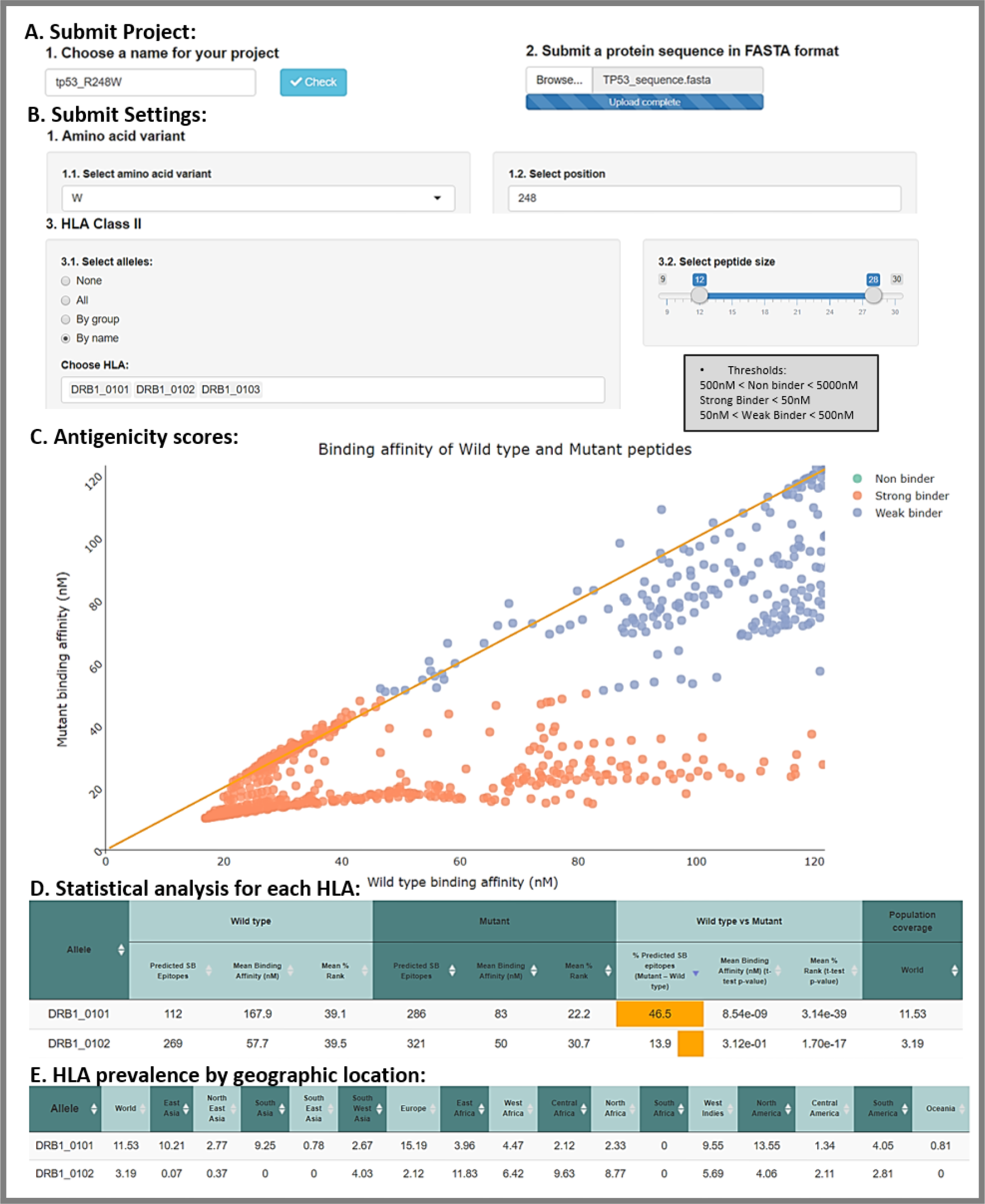
Features of HABIT webserver. A and B) User-definable parameters: protein and amino acid mutation of interest. C) Antigenicity scores: visual comparison of the HLA-binding affinity (nM) between wild-type (x axis) and mutant (y axis) peptides. Classification of peptides in non binder=green; strong binder(SB)=orange and weak binder(WB)=purple. D) Summary and statistical analysis of prediction metrics (binding affinity and % rank) for each HLA allele. E) HLA frequency across 17 world geographic regions.

Overall, HABIT provides a fast and user-friendly way to study neoepitopes, answering the following questions: what is the predicted impact of amino acid mutations in HLA-binding properties?; are the differences statistically significant?; and what percentage/subset of the human population is more likely to respond to the neoepitopes? With this, HABIT might advance the discovery of molecular determinants that influence variation in T cell-mediated immune responses, thus contributing to the design of immune system-based therapy and diagnostic tools.

